# Microglial brain-derived neurotrophic factor (BDNF) supports the behavioral and synaptogenic effects of ketamine

**DOI:** 10.1101/2025.05.05.652266

**Authors:** Samuel C. Woodburn, Alexander Kuhn, David T. Dadodsky, Etienne Mueller, Justin L. Bollinger, Rosa Maria Salazar Gonzalez, J. Elliott Robinson, Lauren Larke Vollmer, Eric S. Wohleb

## Abstract

Microglia have been implicated in the pathogenesis for several psychiatric disorders, yet comparatively little is known about their role in treatments for these conditions. Prior work showed that the rapid-acting antidepressant ketamine increases synaptic density in the prefrontal cortex (PFC), and that brain-derived neurotrophic factor (BDNF) signaling is required for its synaptic and behavioral effects. These studies assumed that neurons were the primary source of BDNF, but other studies have since demonstrated that microglia can produce BDNF in the brain. Still, it remains unclear if microglial BDNF is important for the antidepressant-like effects of ketamine. Our initial studies show that the behavioral and synaptic effects of ketamine are associated with increased *Bdnf* expression in sorted PFC microglia 24 hours after injection. We then demonstrate that conditional BDNF depletion in microglia (*Cx3cr1^Cre/+^:Bdnf^fl/fl^*) reduces GluN2B levels in PFC synaptoneurosomes and attenuates antidepressant-like responses following ketamine treatment compared to genotype controls (*Cx3cr1^Cre/+^:Bdnf^+/+^*). Consistent with this, we found that *Cx3cr1^Cre/+^:Bdnf^fl/fl^* mice show no change in dendritic spine density in the PFC following ketamine. These results indicate that microglial BDNF is important for the effects of ketamine on brain and behavior, expanding upon the role of microglia in pharmacological interventions for psychiatric disorders.

## Introduction

Major depressive disorder (MDD) is a mental health condition characterized by feelings of hopelessness and loss of pleasure (APA, 2013; Kessler et al., 2005; Seedat et al., 2009). Despite the clear impact MDD has on individuals living with it, first-line treatments can take weeks to become effective, with some patients not responding to these treatments at all (Trivedi et al., 2006). Several lines of evidence suggest that low-dose ketamine is effective in treatment-resistant patients, reducing depressive symptoms within 24 hours and lasting for up to two weeks (Berman et al., 2000; DiazGranados et al., 2010; Zarate et al., 2006). Despite these encouraging results, ketamine treatment often requires patients to undergo multiple drug infusions with concerning side effects (Diamond et al., 2014; Rasmussen et al., 2013; Short et al., 2018; Singh et al., 2016). As such, there is a need to identify the cell types and molecular mechanisms underlying the therapeutic properties of ketamine so that more refined treatment options can be developed.

Preclinical studies have uncovered multiple molecular and cellular pathways that contribute to the behavioral and cognitive effects of ketamine. Early work showed that at antidepressant-relevant doses, ketamine increases glutamate signaling in the medial prefrontal cortex (PFC), and additional work shows that this is critical for its rapid effects (Autry et al., 2011; Hare et al., 2020; Homayoun & Moghaddam, 2007; Lepack et al., 2015; Lorrain et al., 2003; Moda-Sava et al., 2019b). Further studies demonstrated that this glutamate burst promotes formation of dendritic spines along the apical branches of PFC pyramidal neurons within 24 hours after ketamine administration (Li et al., 2010b; Moda-Sava et al., 2019b). Importantly, these newly formed spines are required for the lasting behavioral effects of ketamine (Moda-Sava et al., 2019b).

Additional work further showed that ketamine-induced synaptogenesis depends on brain-derived neurotrophic factor (BDNF), as blocking BDNF signaling with antagonizing antibodies limited synaptogenesis and attenuated behavioral effects of ketamine (Lepack et al., 2015). These findings supported other work showing that mice with the Val66Met polymorphism in the BDNF allele, which impairs BDNF release, show baseline deficits in synaptic plasticity and no ketamine-induced changes to brain or behavior 24 hours after administration (Egan et al., 2003; R. J. Liu et al., 2012). However, these studies assumed that neurons were the sole source of BDNF mediating this effect. More recent studies have challenged this assumption, demonstrating that non-neuronal cells express *Bdnf* in the brain (Matt et al., 2018). Microglia have emerged as a potential player in BDNF signaling. Microglia-specific depletion of BDNF was shown to reduce synaptogenesis during learning tasks, resulting in worse performance. (Parkhurst et al., 2013). Our recent studies substantiate these findings as we found that microglial BDNF depletion reduced synaptosome levels of GluN2B and increased susceptibility to the behavioral and synaptic effects of a sub-chronic stress paradigm (Woodburn et al., 2023; H. Yu et al., 2012). These studies indicate that microglial BDNF supports adaptive neuroplasticity and notably influences synaptic GluN2B levels which is important for the effects of ketamine (Gerhard et al., 2020a; Y. Zhang et al., 2021a).

In this context, we carried out studies to determine if microglial BDNF is important for the synaptic and behavioral effects of ketamine. This work showed that the behavioral and synaptogenic effects of ketamine are associated with a two-fold increase in microglial *Bdnf* expression 24 hours post-ketamine treatment. Further studies showed that conditional BDNF depletion in microglia (*Cx3cr1^Cre/+^:Bdnf^fl/fl^*) decreased GluN2B expression in the medial PFC, and limited the behavioral and synaptogenic effects of ketamine. These studies provide initial evidence that microglial BDNF is important for the neurobiological effects of low-dose ketamine.

## Methods and Materials

### Animals

Studies were performed using adult male mice (7-8 weeks old). Mice were group-housed (4/cage) in 11.5”x 7.5”x 6” polypropylene cages under a 12 h light-dark cycle (light on: 08:00-20:00) with *ad libitum* access to water and rodent chow. Thy1-GFP-M mice were obtained from in-house breeders (Jackson Laboratories; Tg(Thy1-EGFP)MJrs/J; #007788). Wild-type C57BL/6 mice were purchased from Jackson Laboratories (#000664). Breeder pairs for the *Cx3cr1^CreER^* (*B6.129P2(Cg)-Cx3cr1^tm2.1(cre/ERT2)Litt/WganJ^*; #021160) and *Bdnf^2lox^* (*Bdnf^tm3Jae^/J*; #004339) strains were purchased from Jackson Laboratories and maintained in-house (Parkhurst et al., 2013; Rios et al., 2001). The mouse line was maintained as being heterozygous for the *Cx3cr1^CreER^* allele and homozygous for either the *Bdnf^+/+^* or *Bdnf^fl/fl^*alleles. All breeder pairs for mice used in this study were generated from the F1 generation of a cross between *Cx3cr1^CreER/CreER^*and *Bdnf^fl/fl^* mice. Animal experiments were carried out in accordance with the National Institutes of Health guide for the care and use of laboratory animals and studies were approved by the University of Cincinnati Institutional Animal Care and Use Committee. As such, all efforts were made to minimize animal suffering and reduce the number of animals used.

### Tamoxifen administration

Mice were placed on a tamoxifen diet for 3 weeks (80 mg/kg; Envigo, #TD.130585) after weaning on postnatal day 21-25. After these 3 weeks, tamoxifen diet was replaced with standard vivarium diet (Teklad LM-485 diet, 7912). Mice received vivarium diet for 2 weeks prior to ketamine treatment to allow peripheral *Cx3cr1^CreERT2/+^* expressing cells to turnover from bone marrow progenitors while microglia populations remained relatively stable (Harley et al., 2021; Parkhurst et al., 2013; Woodburn et al., 2023).

### Ketamine administration

Mice received a single injection of ketamine hydrochloride (Zoetis) at 10 mg/kg body weight or vehicle solution (0.9% NaCl). Drugs were injected intraperitoneally (i.p.) with a volume of 10 mL/kg body weight. Dosing was based on prior literature (Li et al., 2010b).

### AAV-PHP.eB-hSyn1-tdTomato Administration

Following viral production and titering, AAV vectors were intravenously administered via injection into the retro-orbital sinus with an insulin syringe (BD U-100 Ultrafine 0.5 mL syringe with 31-gauge 6mm needle) during anesthesia with isoflurane (1-3% in 95% O_2_/5% CO_2_ provided via nose cone at 1 L/min) as previously described (Challis et al., 2019). Following the r.o. injection, mice were allowed to recover in a designated biocontainment room

### Behavioral testing

Forced swim test (FST) was conducted as previously described. (Woodburn et al., 2021a). All behavioral tests were performed in a normally lit room (300 lux, white light), during the light phase of the circadian cycle (08:00-11:00). Mice were placed in the room 30 minutes prior to testing to habituate. For the FST, mice were placed in a 2-liter beaker of water (24°+/-1°C) for 8 minutes and time spent immobile was determined (Can et al., 2011).

### Percoll gradient enrichment of microglia

Brain microglia were enriched as previously described (Horchar & Wohleb, 2019). Dissected frontal cortex was passed through a 70 µm cell strainer. Homogenates were centrifuged at 1200×g for 5 min. Supernatants were removed and cell pellets were re-suspended in 70% isotonic Percoll (GE Healthcare, Uppsalla, Sweden, #17089102). A discontinuous Percoll density gradient was layered as follows: 30% and 0% isotonic Percoll. The gradient was centrifuged for 20 min at 2000×g and enriched microglia were collected from the interphase between the 70% and 30% Percoll layers. Enriched microglia were labeled with antibodies for flow cytometry and sorted based on CD11b/CD45 expression using a BioRad S3e cytometer/cell sorter (Hercules, CA, U.S.A.).

### Fluorescence-activated cell sorting (FACS)

Staining of cell surface antigens was performed as previously described (Horchar & Wohleb, 2019). In brief, Fc receptors were blocked with anti-CD16/CD32 antibody (BioLegend, San Diego, CA, U.S.A., #553141). Cells were washed and then incubated with conjugated antibodies (FITC-CD115, #135512 Biolegend; PE-P2Y12, #848004 Biolegend; PE-CF594-CD45, #562420 BD Biosciences; PerCp-Cy5.5-CD11b, #550993 BD Biosciences) for 1 h at 4^ο^C. Cells were washed and then re-suspended in FACS buffer for analysis. Non-specific binding was assessed using isotype-matched antibodies. Antigen expression was determined using a BioRad S3e four-color cytometer/cell sorter. Data were analyzed using FlowJo software (Ashland, OR, U.S.A.).

### RNA isolation and quantitative real-time PCR

RNA was extracted from microglia using a Single Cell RNA purification kit (Norgen Biotek, Thorold, Canada, #51800). Samples were reverse transcribed and quantitative real-time PCR was conducted as previously described (Horchar & Wohleb, 2019).

### Immunohistology

Mice were transcardially perfused with sterile PBS and 4% paraformaldehyde (PFA). Brains were post-fixed in 4% PFA for 24 h and incubated in 30% sucrose for an additional 24 h. Frozen brains were sectioned at 40 µm using a Leica CM3050 S cryostat. Free-floating sections were washed, then blocked for 1 hour at room temperature. Sections were washed, then incubated with rabbit anti-IBA1 (Wako, #019-19741) overnight at 4°C. Sections were washed and incubated with conjugated secondary antibody overnight at 4°C (Alexa Fluor 546 donkey anti-rabbit). Following secondary antibody incubation, sections were washed, mounted on Superfrost(+) slides (Fisher Scientific) with Fluoromount-G, and coverslipped (Fisher Scientific).

### Quantitative immunofluorescence

Confocal images were obtained on a Nikon C2^+^ microscope equipped with CFI Plan Apochromat Lambda Series Objectives. Images were captured with Nikon C2si^+^ camera and analyzed using Nikon Elements software. In Thy1-GFP(M) mice, layer I of the medial PFC was identified and apical dendritic segments were imaged using 60x objective lens with a 2.5x zoom (NA 1.4) with z-stack sampling: 0.075 µm. For each sample, 6-8 dendritic segments were analyzed using Filament Tracer in Imaris. Proportional area of IBA1^+^ material was measured using standardized threshold parameters with ImageJ software (NIH, Bethesda, Maryland). IBA1^+^ cells were counted by a trained researcher blinded to animal condition (4-6 images per sample).

### Isolation of synaptoneurosome fractions

Synaptoneurosomes were isolated using previously published methods (Li et al., 2010b). A collection buffer was prepared the evening prior to sacrifice by combining the following reagents: 320 mM sucrose, 0.5 mM CaCl_2_, 1 mM NaCHO_3_, 0.1 mM PMSF, 4 mM HEPES buffer (Fisher Scientific, #BP299-100), and one cOmplete^™^ EDTA-free protease inhibitor cocktail tablet (Roche diagnostics, #11836170001). Mice were sacrificed via cervical dislocation 2 hours after the final stressor (day 8), at which point the medial PFC was dissected and homogenized in 1 mL of collection buffer via 12 up and down strokes of pestle B of a 7 mL Dounce homogenizer. Samples were then fractionated by centrifugation at 800 × g for 10 min. at 4 °C, yielding a pellet containing the nuclear fraction (P1) and a protein-rich supernatant (S1). The S1 fraction was collected into a new tube and centrifuged again at 13,800 × g (12,000 rpm) for 10 min at 4 °C to yield a pellet containing crude synaptoneurosomes (P2) and a cytosolic fraction as supernatant (S2). The S2 fraction was pipetted into a new tube, and all samples were stored at -80 °C until use.

### Western blotting

P2 fractions of synaptoneurosomes were resuspended in 250 µL of RIPA lysis buffer (50 mM Tris HCl pH 7.4, 150 mM NaCl, 1 mM NaVO_3_, 10 mM NaF, 0.1% SDS, 5% Triton X-100, and a cOmplete^™^ EDTA-free protease inhibitor cocktail tablet). Protein content was determined using a BCA assay (ThermoFisher #23227). 20 µg of each sample was combined with dH_2_O, 7.5 µL NuPAGE^™^ 4× LDS buffer (ThermoFisher Scientific, #NP0007) and 3 µL β-mercaptoethanol for a volume of 30 µL, then denatured at 95 °C for 5 minutes. Samples were then loaded into a 4-12% Bis-Tris gel (GenScript, #M00654) along with SeeBlue Plus2 and Magic Mark XP protein standards (ThermoFisher Scientific, #LC5925; #LC5602). Electrophoresis was conducted at 200 V with Tris-MOPS-SDS running buffer (GenScript, #M00138) in an XCell SureLock mini-cell system (ThermoFisher Scientific, #EI0002) with a PowerPac basic (Bio-Rad, #1645050). Protein was then transferred onto a 0.45 μm pore-size PVDF membrane (Cytiva, #45-004) with a 10% methanol transfer buffer (GenScript, #M00139) at 30 V constant for 1 hour using an XCell II Blot Module (ThermoFisher, #EI0002). No-Stain^™^ Protein Labeling Reagent (ThermoFisher Scientific, #A44717) was applied after transfer per the manufacturer’s protocol. Membranes were then washed and blocked for 1 hour with 5% BSA + 1× TBST. Primary antibodies for GluA1 (1:5,000; Antibodies Inc., #N355/1), GluA2 (1:2,000: Antibodies Inc., #L21/32), GluN2A (1:500; Antibodies Inc., #N327/95), and GluN2B (1:500; Antibodies Inc., #N59/20) were diluted in 5% BSA + 1× TBST and incubated with membranes overnight at 4 °C. The next day, membranes were washed and incubated at room temperature for one hour in donkey anti-mouse HRP conjugated secondary antibody (1:10,000; Jackson Immuno Research, #715-035-150) in 5% BSA + 1× TBST. Afterward, membranes were washed with 1× TBST, and then ECL reagents (ThermoFisher Scientific, #32106) were applied for 5 minutes. Imaging for total protein and ECL was conducted using an iBright CL750 (Thermofisher Scientific, #A44116) and image analysis was conducted using iBright analysis software. Integrated density for proteins of interest was normalized by the total protein content within the same lane (Aldridge et al., 2009).

### Statistics

Data were analyzed using GraphPad Prism 9.3.1 (La Jolla, California). Outliers were removed using Grubb’s test (α = 0.05). Comparisons made between wildtype C57BL/6 or Thy1-GFP (M) mice given ketamine or vehicle were made using unpaired t-tests. Effects of ketamine or vehicle on *Cx3cr1^CreER/+^:Bdnf^+/+^*or *Cx3cr1^CreER/+^:Bdnf^fl/fl^*mice were determined using a 2 × 2 ANOVA (genotype × drug), with a Holm-Šidák’s multiple comparison test used to assess between group differences.

## Results

### Synaptogenic and behavioral effects of ketamine do not correspond to changes in microglial morphology

To confirm that ketamine exerts synaptogenic and behavioral effects in our hands, male Thy1-GFP (M) mice were treated with vehicle or ketamine (10 mg/kg). Immobility in the FST was then assessed 24 hours following injections. Within 2 hours of behavioral testing, mice were transcardially perfused for immunohistology (Fig.1A). Analysis of behavioral data showed that ketamine decreased immobility in the FST compared to vehicle-treated controls (t(9) = 3.04; *p* = 0.014; Fig.1B). Thy1-GFP (M) mice express GFP in a subset of pyramidal neurons, allowing us to measure apical spine density in layer I of the medial PFC. Similar to previous reports, we found that ketamine treatment increased apical dendritic spine density in the medial PFC compared to vehicle-treated controls (t(7) = 3.885; *p* = 0.006; Fig.1C-D) (Li et al., 2010b). Additional sections were immunolabeled for IBA1 to measure effects of ketamine on microglial morphology (Fig.1E). However, we did not find any significant differences in the number or proportional area of IBA1+ cells (Fig.1F-G), but there was a modest decrease in the nearest neighbor distance (NND, t(11) = 2.663; *p* = 0.022; Fig.1H).

**Figure 1.**
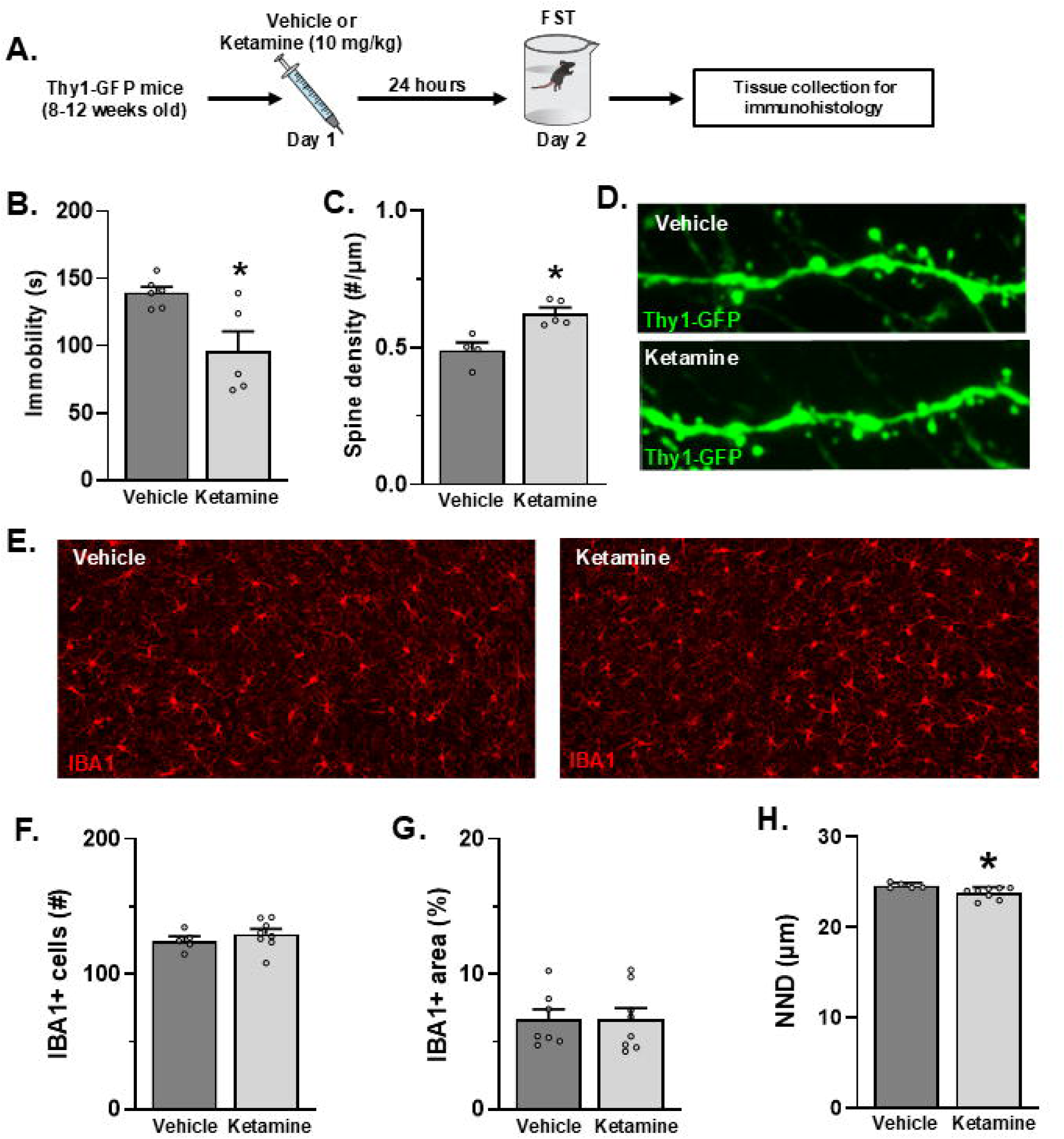
Behavioral and synaptic effects of ketamine do not correspond to morphological changes in PFC microglia. Male Thy1-GFP(M) mice were treated with vehicle or ketamine (10 mg/kg). Behavioral testing was conducted 24 hours later, followed by tissue collection for immunohistology. **A)** Schematic showing experimental approach and timeline. **B)** Immobility in the FST was assessed (*n* = 5-6/group). **C)** Images of apical dendritic spines in layer I of the medial PFC were acquired and **(D)** density of spines was assessed (*n* = 4-5/group). **E)** Representative images of IBA-1^+^ microglia in the medial PFC were collected. **F)** The number (*n* = 5-7/group) and **(G)** proportional area (*n* = 8/group) of microglia was quantified. Bars represent the mean ± S.E.M. Means significantly different than vehicle-treated control animals are denoted (*, *p* < 0.05) based on unpaired t-test results.

### Microglial Bdnf is upregulated 24 hours after ketamine administration

Previous work indicates that ketamine increases *Bdnf* transcription, but the specific contributions of different cell types remain unclear (Autry et al., 2011). To determine if microglia are involved in this effect, we treated male C57BL/6 mice with ketamine (10 mg/kg) or vehicle as control. After 24 hours, mice were rapidly euthanized via cervical dislocation, and samples of frontal cortex were collected and microglia were isolated via FACS (Fig.2A-C). Analysis of MFI data collected during flow cytometry did not indicate any changes in microglial surface expression of CSF1R, P2Y12, or CD11b (Fig.2D). RNA was isolated from sorted microglia samples, and transcriptional changes were assessed via qPCR. We found that ketamine caused a slight increase in *Csf1r* expression (t(14 )= 2.625; *p* = 0.02) while *Bdnf* transcript expression was increased by about two-fold (t(14) = 3.243; *p* = 0.007; Fig.2D).

**Figure 2.**
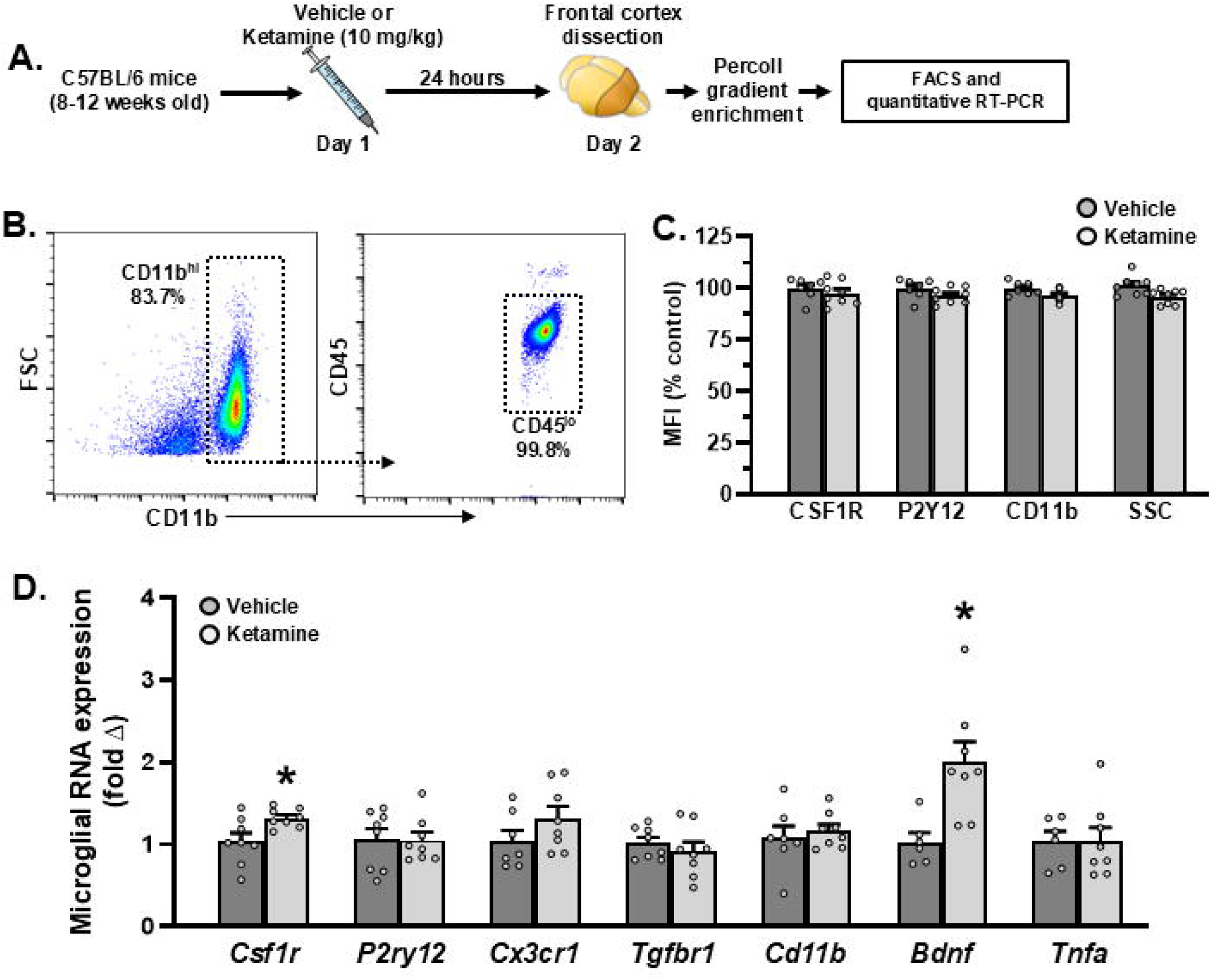
Ketamine increases microglial *Bdnf* expression in the frontal cortex. Male C57BL/6 mice were treated with vehicle or ketamine (10 mg/kg). Approximately 24 hours later, frontal cortex samples were collected and processed for FACS. **A)** Schematic showing experimental approach and timeline. **B)** Gating strategy for isolation of CD11b^+^, CD45^lo^ microglia is shown in the representative dot plot. **C)** The relative purity of sorted microglia samples is shown (*n* = 7-8/group). **D)** Expression of cell surface markers in sorted microglia was assessed using MFI data (*n* = 7-8/group). Following FACS, qPCR was performed on isolated microglia samples. **E)** Relative expression of *Bdnf* is shown (*n* = 10-12/group). Bars represent the mean ± S.E.M. Means significantly different than vehicle-treated control animals are denoted (*, *p* < 0.05) based on unpaired t-test results.

### Depletion of microglial BDNF abolishes ketamine’s effect in the FST

To test the importance of microglial BDNF signaling to ketamine’s synaptic and behavioral effects, we utilized transgenic mice with a tamoxifen-inducible Cre recombinase under the monocyte specific *Cx3cr1* promoter (Parkhurst et al., 2013). In these experiments, male *Cx3cr1^CreER/+^:Bdnf^+/+^* and *Cx3cr1^CreER/+^:Bdnf^fl/fl^*mice were supplied with tamoxifen diet for 21 days. After this, tamoxifen diet was switched out for standard vivarium chow for 14 days. This strategy leverages the long lifespan of microglia compared to peripheral *Cx3cr1^+^*cells, which replenish from *Cx3cr1^−^* progenitors, thereby limiting recombination to microglia 14 days after tamoxifen administration has ceased (Parkhurst et al., 2013; Woodburn et al., 2023). Following this, mice of both genotypes were randomly assigned to receive ketamine (10 mg/kg) or vehicle as control. Mice were subjected to the FST 24 hours post-injection, then sacrificed 30 minutes later. Synaptoneurosomes were isolated from samples of the medial PFC immediately following this for later quantification of protein expression via western blotting (Fig.3A).

**Figure 3.**
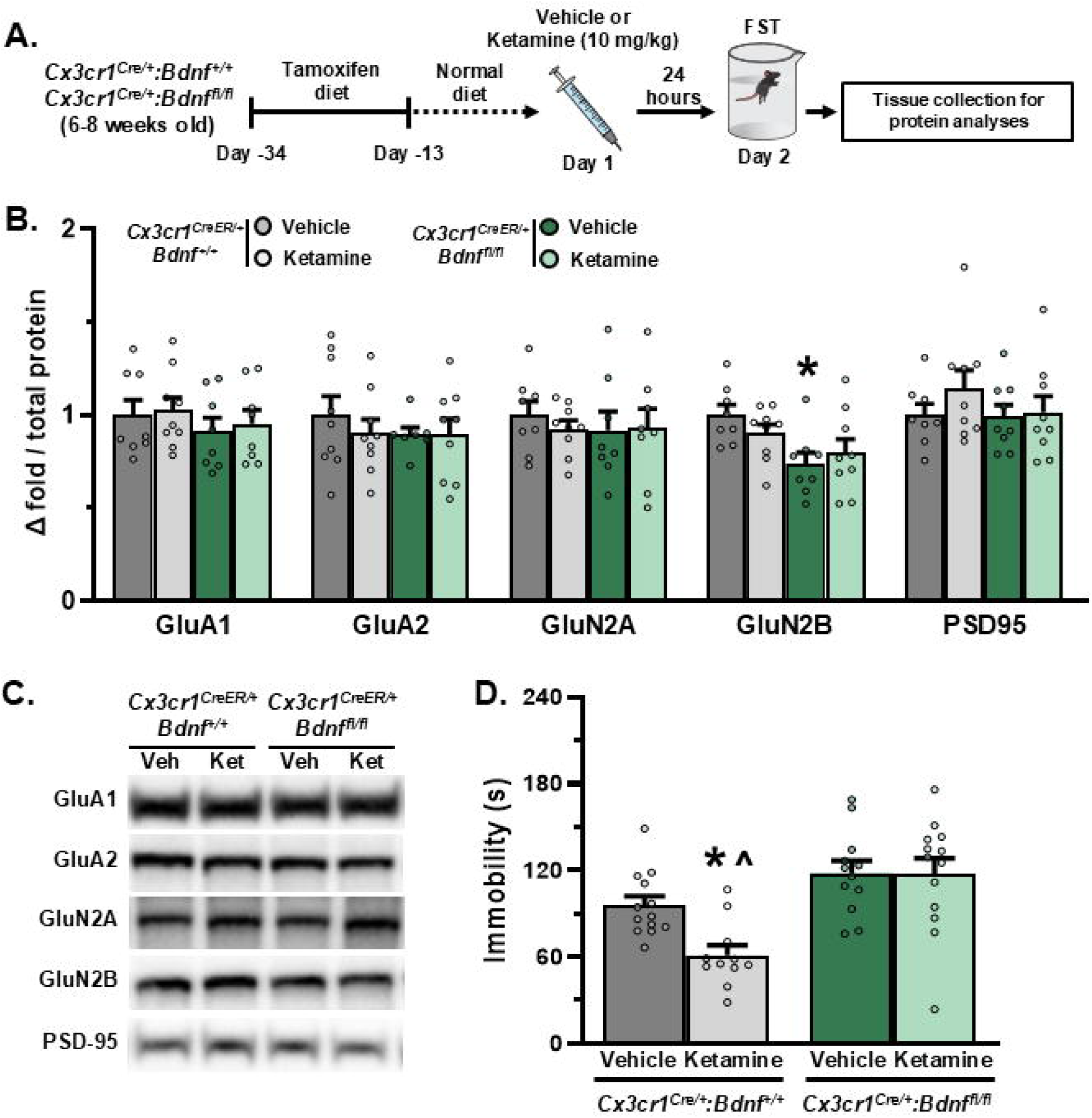
Microglial BDNF depletion abrogates ketamine’s effect in the forced swim. Male *Cx3cr1^CreER/+^:Bdnf^+/+^*and *Cx3cr1^CreER/+^:Bdnf^fl/fl^* mice were supplied with a tamoxifen diet for 21 days to induce Cre-mediated recombination. After this, 14 days were given for peripheral *Cx3cr1^+^* cells to replenish. Mice were then treated with vehicle or ketamine (10 mg/kg). After 24 hours, behavior was assessed in the FST, and medial PFC samples were collected for synaptosome isolation. **A)** Schematic showing experimental approach and timeline. Protein expression was assessed in medial PFC synaptosomes via western blot. **B)** Quantification of GluA1, GluA2, GluN2A, GluN2B, and PSD-95 expression (*n* = 7-9/group) relative to total lane protein (see Fig S1). **C**) Representative bands for synaptic proteins are shown. **D)** Immobility in the FST was assessed (*n* = 11-13/group). Statistical significance was determined via 2 × 2 ANOVA followed by Holm-Šidák’s multiple comparison test. Bars represent the mean ± S.E.M. Based on post-hoc analysis, means significantly different than controls of the same genotype are denoted (*, *p* < 0.05), and means significantly different from opposite genotype controls are denoted (^, *p* < 0.05).

Analysis of synaptic proteins revealed that *Cx3cr1^CreER/+^:Bdnf^fl/fl^* mice displayed a baseline deficit in GluN2B expression (genotype: F_1,30_ = 9.247; *p* = 0.0049), which was most pronounced compared to vehicle-treated *Cx3cr1^CreER/+^:Bdnf^+/+^* mice (*p* = 0.0339; Fig.3B-C and S1). However, none of the other proteins analyzed differed significantly due to ketamine treatment or microglial BDNF depletion. Nonetheless, assessment of FST data showed that, while ketamine decreased immobility for *Cx3cr1^CreER/+^:Bdnf^+/+^* mice (interaction: F_1,45_ = 4.1; *p* = 0.048; Fig.3D), ketamine-treated *Cx3cr1^CreER/+^:Bdnf^fl/fl^* mice showed no difference in FST immobility from vehicle-treated controls (*p* = 0.962).

### Synaptogenic effects of ketamine are not observed in mice lacking microglial BDNF

Seminal studies have demonstrated that the behavioral effects of ketamine are dependent on synapse formation in the medial PFC, resulting in increased dendritic spine density on the apical dendrites of pyramidal neurons (Li et al., 2010a; Moda-Sava et al., 2019a). To examine changes in dendritic spine density we utilized a systemic AAV approach to sparsely label neurons in the cortex. The AAV-PhP.eB serotype is effective at crossing the blood-brain barrier and limits the need for intracranial surgery (Challis et al., 2019). Male *Cx3cr1^CreER/+^:Bdnf^+/+^* and *Cx3cr1^CreER/+^:Bdnf^fl/fl^*mice were supplied with a tamoxifen diet for 21 days. After this, mice received a retroorbital injection of AAV-PhP.eB-hSyn1-tdTomato and were switched to standard vivarium chow for 14 days. This allowed for viral gene expression and for turnover of peripheral *Cx3cr1^+^* cells. Mice of both genotypes were randomly assigned to receive ketamine (10 mg/kg) or vehicle as control, and then 24 hours later brain were collected for immunohistology (Fig.4A&B). Semi-automated reconstruction of dendritic spine segments showed that ketamine significantly increased spine density in *Cx3cr1^CreER/+^:Bdnf^+/+^* (*p* < 0.007 compared to all groups), but spine density in *Cx3cr1^CreER/+^:Bdnf^fl/fl^* was not changed (interaction: F_1,19_ = 9.243; *p* = 0.0067; Fig.4C&D).

**Figure 4.**
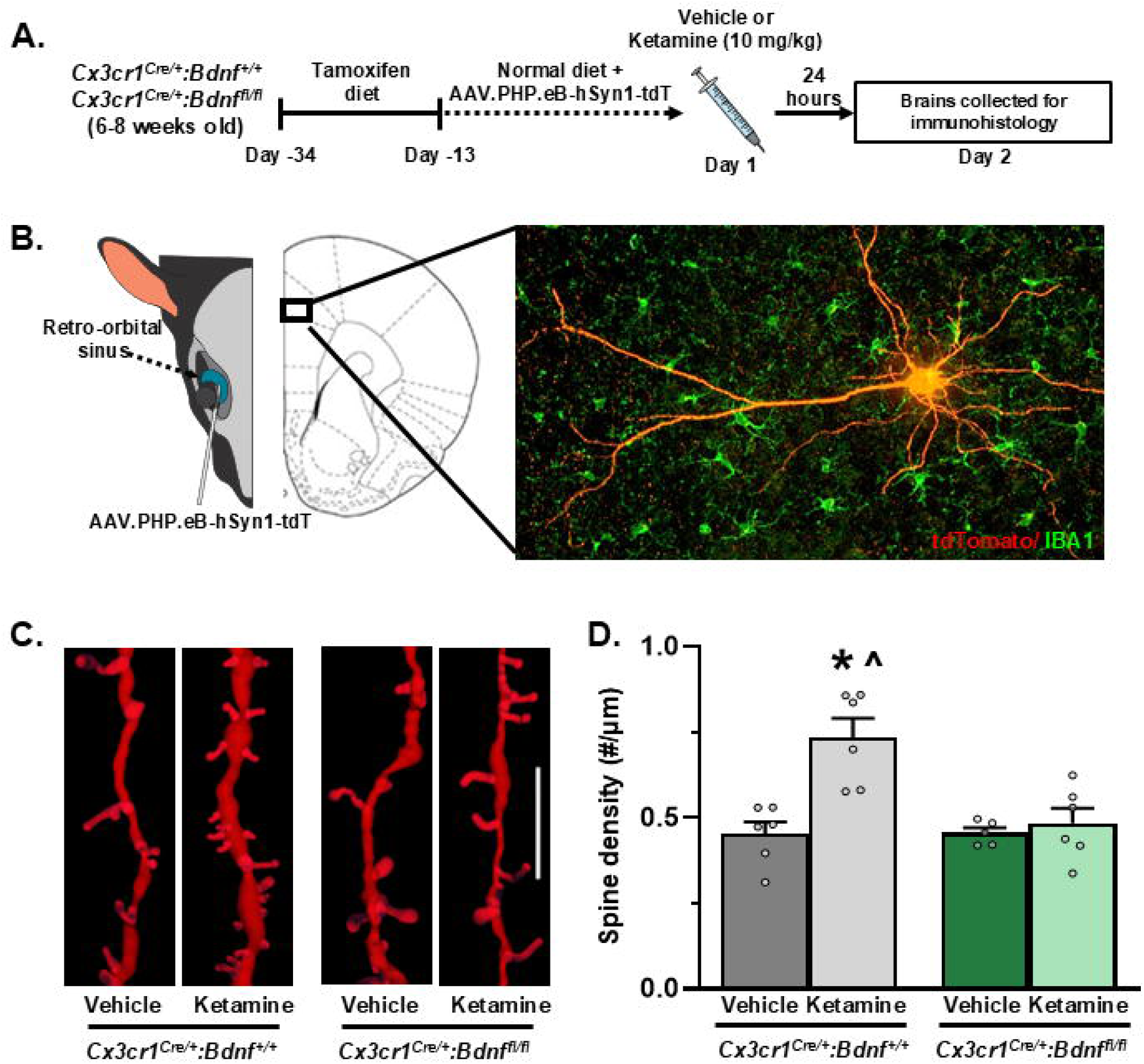
Synaptogenic effects of ketamine are not observed in mice lacking microglial BDNF. Male *Cx3cr1^CreER/+^:Bdnf^+/+^*and *Cx3cr1^CreER/+^:Bdnf^fl/fl^* mice were supplied with a tamoxifen diet for 21 days to induce Cre-mediated recombination. After this, mice received a retroorbital injection of AAV.PhP.eB-hSyn1-tdTomato to induce sparse labeling of neurons in the brain. Mice were given 14 days to allow viral expression and peripheral *Cx3cr1^+^* cells to replenish. Mice were then treated with vehicle or ketamine (10 mg/kg). After 24 hours, mice were perfused and brains collected for immunohistology. **A)** Schematic showing experimental approach and timeline. **B)** Cartoon depicting retroorbital AAV injection and representative viral expression in PFC pyramidal neuron with immunolabeling for IBA1+ cells. **C)** Representative images of dendritic segments in layer I of the medial PFC. Scale bar represents 10 µm. **D)** Average spine density is shown. Statistical significance was determined via 2 × 2 ANOVA followed by Holm-Šidák’s multiple comparison test. Bars represent the mean ± S.E.M. Based on post-hoc analysis, means significantly different than controls of the same genotype are denoted (*, *p* < 0.05), and means significantly different from opposite genotype controls are denoted (^, *p* < 0.05).

## Discussion

Preclinical research over the past two decades have demonstrated that the lasting behavioral effects of ketamine are due to novel spine formation in corticolimbic brain regions such as the medial PFC, and that this depends on BDNF signaling (Autry et al., 2011; Kim et al., 2021; Lepack et al., 2015; Li et al., 2010a; R. J. Liu et al., 2012; Moda-Sava et al., 2019b). Indeed, mice with the Val66Met BDNF mutation, which impairs BDNF release, display reduced spine formation at baseline, and ketamine has minimal effects on synaptic plasticity or behavior in these mice (Egan et al., 2003; R. J. Liu et al., 2012). Additionally, administration of BDNF neutralizing antibodies in the medial PFC is sufficient to replicate this effect (Lepack et al., 2015). It was broadly assumed that BDNF was only released from synaptic compartments, and thus these studies did not consider the contributions of non-neuronal cells in this signaling pathway (Dieni et al., 2012; Hartmann et al., 2001; Zakharenko et al., 2003). However, more recent data suggests that many cell types express *Bdnf* in the brain, and depletion of microglial BDNF in particular has been linked to deficits in synaptic plasticity (Matt et al., 2018; Parkhurst et al., 2013). Here, we show that, while ketamine does not obviously shift the morphology or surface marker expression of PFC microglia, it does increase microglial *Bdnf* expression. Further, we show that mice with deficient microglial BDNF signaling show limited synaptic and behavioral effects at 24 hours after ketamine administration. Collectively, these studies add to a growing literature suggesting that microglia are important for the effects of antidepressant treatments (Kreisel et al., 2014; Rimmerman et al., 2021; J. Zhang et al., 2021).

Historically, much of the work characterizing microglial function has focused on their role in injury, illness, and disease. In these conditions, there are often profound changes in microglial phenotype and function which are associated with neurological and behavioral outcomes of the disease model being studied (Estes & McAllister, 2014; Kierdorf & Prinz, 2013; Woodburn et al., 2021b). This has led some to classify microglia into discrete phenotypic categories, with the underlying assumption that these changes will predict specific brain pathologies (Butovsky et al., 2013; Keren-Shaul et al., 2017; Stratoulias et al., 2019; Zrzavy et al., 2017). While evidence suggests this may be the case in neuroinflammatory conditions like multiple sclerosis or Alzheimer’s disease, our results indicate that not all biologically relevant changes in microglia present obvious changes in phenotype. Indeed, had we limited our characterization of microglia to morphology and cell surface receptor composition, it might have appeared that ketamine did not affect microglia at all. Further, it should be noted that compiling evidence suggests the signaling pathways implicated in injury and disease can have drastically different functions under homeostatic conditions of (Garré et al., 2017; Goshen et al., 2007; Song et al., 2013). Indeed microglial sources of BDNF have historically been studied for their function in pathology, such as the synaptic potentiation underlying chronic pain (Coull et al., 2005; Ulmann et al., 2008; Zhou et al., 2019). This underscores the fact that microglial functions outside of overtly inflammatory conditions are still incompletely understood and cautions against limiting investigations of microglia to markers associated with their immune functions.

Microglial morphology in the brain is highly dynamic and shifts in response to activity-dependent neuronal cues (Davalos et al., 2005; Dissing-Olesen et al., 2014; Eyo et al., 2014, 2015a; Y. U. Liu et al., 2019; Stowell et al., 2019). It is also well-established that ketamine drastically increases excitatory transmission in the medial PFC within the first few hours of administration, only to return to physiological levels soon after (Hare et al., 2020; Lorrain et al., 2003; Moda-Sava et al., 2019b). Thus, it is possible that microglia do show obvious changes in morphology or neuronal surveillance in the initial period after ketamine administration, but these changes are lost 24 hours after ketamine administration. Future studies using *in vivo* imaging may reveal how ketamine influences interactions between microglia and the cortical neurons. Considering that neuron-microglia interactions are highly influenced by fluctuations in dendritic calcium signaling and that ketamine increases calcium signaling from dendritic spines in the frontal cortex, such studies may prove particularly fruitful (Ali et al., 2020a; Eyo et al., 2015b, 2018).

Findings from chronic pain models may also have a great deal of relevance here, as microglial BDNF has consistently been associated with increased neuronal activity in the dorsal horn after spinal injury. Specifically, increased glutamatergic signaling from spinal neurons leads to neuronal release of activity-dependent factors, such as purines (ATP/ADP) and CSF1 (Trang et al., 2009; Zhou et al., 2019). Microglia detect these signals via purine receptors (i.e. P2X4, P2X7, P2Y12), and CSF1R, which ultimately leads to the release of BDNF, and long-term potentiation in the dorsal horn (Gong et al., 2009; Kobayashi et al., 2008; Sperlágh & Illes, 2014; Tozaki-Saitoh et al., 2008; Trang et al., 2009; Ulmann et al., 2008; T. Yu et al., 2019; Zhou et al., 2019). Given that microglia in the brain are also sensitive to these activity-dependent factors, it is possible that ketamine increases microglial *Bdnf* expression via these same mechanisms. However, microglia show a great degree of regional heterogeneity (Hickman et al., 2013; Tan et al., 2020). Thus, future studies are needed to determine if similar molecular pathways regulate spinal and cortical microglia.

Our results in microglial BDNF depleted mice confirm earlier findings that microglial BDNF regulates GluN2B expression under baseline conditions (Parkhurst et al., 2013; Woodburn et al., 2023). Importantly, ketamine is known to preferentially antagonize active GluN2B containing NMDA receptors, which at sub-anesthetic doses drives glutamate signaling in the PFC (Homayoun & Moghaddam, 2007; Lorrain et al., 2003; Y. Zhang et al., 2021b). This might provide an explanation for why *Cx3cr1^CreER/+^:Bdnf^fl/fl^*mice did not show the reduced FST immobility and increased spine density typically observed with ketamine administration. It is possible that reduced synaptic GluN2B limits the receptors for ketamine to bind to and thereby diminishes its effects. Additional testing is required to know for certain, yet it is worth considering which neurons are directly affected by microglial BDNF depletion. For the past decade, it has been debated whether the effects of ketamine are due to antagonism of inhibitory interneurons, which express high levels of GluN2B in the PFC, or if they are driven by inhibiting extrasynaptic GluN2B containing NMDA receptors on pyramidal neurons (Abdallah et al., 2018; Monteggia & Zarate, 2015). Evidence supports both sides of this debate, as separate reports indicate GluN2B knockdown in excitatory or inhibitory cells affects the efficacy of ketamine (Ali et al., 2020b; Gerhard et al., 2020b; Miller et al., 2014; Pothula et al., 2021). It is unclear from our results whether microglial BDNF regulates GluN2B expression on excitatory or inhibitory neurons, however further studies can test this compelling question.

There are a few caveats to the present study, however. The first being that we did not observe increased expression of synaptic markers in ketamine-treated *Cx3cr1^CreER/+^:Bdnf^+/+^* mice, despite decreased immobility in the FST and increased dendritic spine density. Prior work reported synaptic GluR1 and PSD-95 were increased in ketamine-treated animals (Li et al., 2010b). It is possible that different methods of protein quantification contributed to conflicting results. Second, while this and prior work suggest that microglial BDNF regulates cortical GluN2B expression, additional studies are needed to test specific mechanisms. It is also worth noting that our protein analyses were limited to glutamate receptors in the PFC. It is well-established that ketamine affects multiple brain regions and transmitter systems (Duman et al., 2016; Witkin et al., 2016; Wu et al., 2021; Yao et al., 2017). Thus, there may be alternative mechanisms to explain why microglial BDNF depletion affects responses to ketamine. In either case, future studies will be needed to better understand the role microglial BDNF plays in supporting the effects of ketamine on the brain and behavior. Last, we limited our analyses to male mice, as our previous work suggested more dynamic regulation of microglial BDNF in males (Woodburn et al., 2021a). The reason for these sex differences is unclear, but other labs have shown sex-dependent effects with ketamine treatment, and microglia appear to regulate *Bdnf* differently between sexes (Carrier & Kabbaj, 2013; Franceschelli et al., 2015; Sorge et al., 2015; Woodburn et al., 2021a). While these baseline effects of sex lie outside the scope of the present study, understanding this will be necessary to understand the full importance of neurotrophic signaling from microglia.

In summary, these experiments suggest that ketamine increases *Bdnf* expression in microglia from the frontal cortex without driving obvious phenotypic changes in morphology or surface receptor expression. Further, we show that depletion of microglial BDNF reduces GluN2B in PFC synaptosomes and limits the behavioral and synaptogenic actions of ketamine. Collectively, this work suggests that microglia is an important mediator of cortical neuroplasticity and in turn the behavioral effects of ketamine.

---

**Figure S1.**
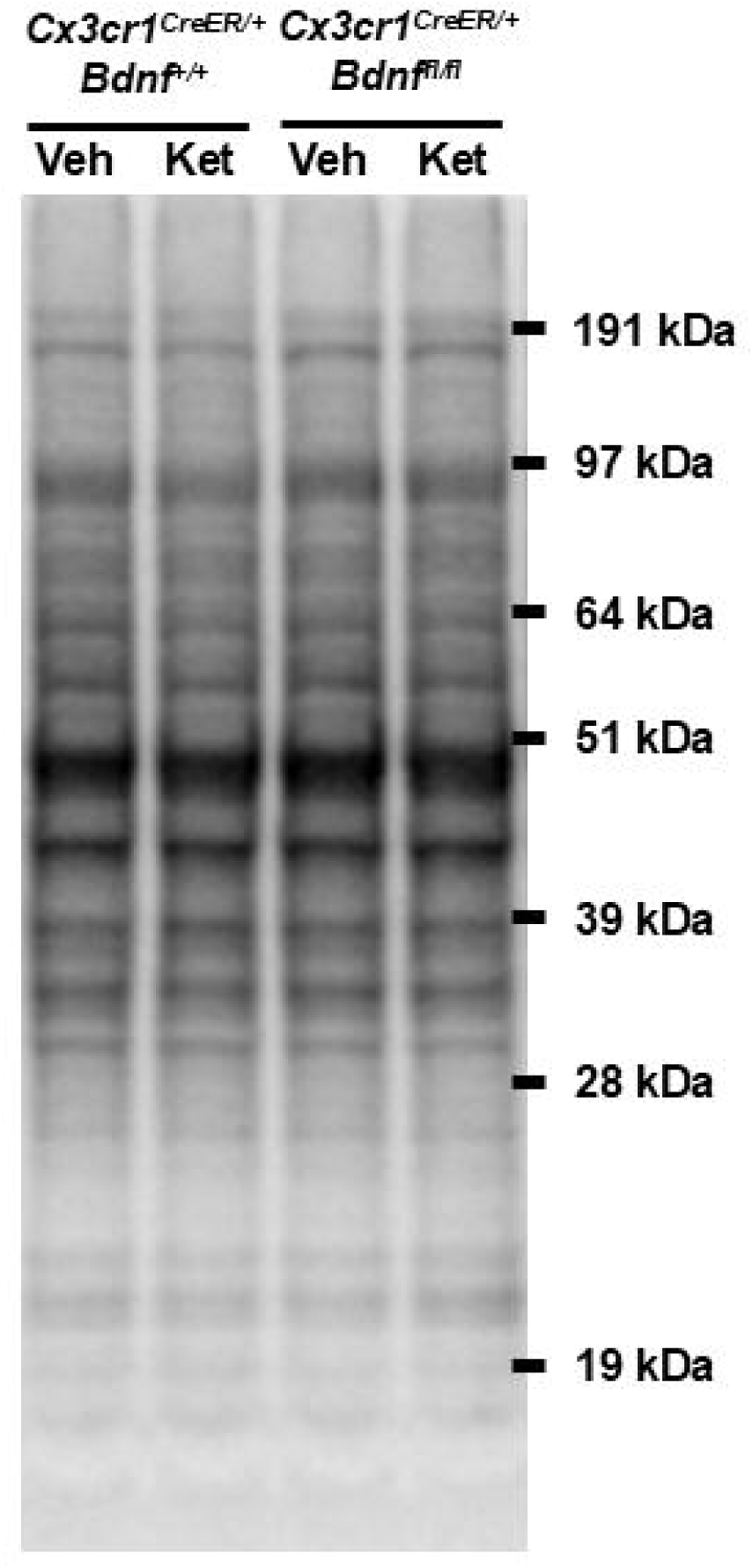
Representative total protein staining. 20 µg of protein were run in each lane of an SDS-Page gel. To calculate the relative density of bands in western blotting, ECL signal was normalized to total lane protein, shown here.

